# Joint Spherical-Harmonics Regression for PheWAS: Global Maps, Residual Localization, and Spherical-Cap Enrichment

**DOI:** 10.1101/2025.10.27.684843

**Authors:** Manuel A. Rivas

**Affiliations:** Department of Biomedical Data Science, Stanford University, Stanford, CA, USA

## Abstract

We present an application for analyzing variant-by-phenotype association summaries from PheWAS [10] using spherical harmonics (SH) [5, 4]. The method embeds phenotypes onto the unit sphere, fits a *joint* SH map across all variants via weighted ridge regression [3, 8, 9] with a Laplacian penalty, and then assesses *per-variant residual* localization beyond this global structure. Rotation-invariant descriptors (degree power spectrum, *l*_95_, entropy, and a localization index) summarize spatial complexity. We detect localized genetic effects with a sign-aware spherical-cap enrichment test that scans cap radii around SH extrema and identifies phenotypes driving hotspots via inverse-variance fixed-effect meta-analysis [7]. Model selection uses BIC [2], and significance for high-degree structure uses a nested *F*-test within the linear-model framework [8, 9] with BH-FDR across variants [1]. The implementation provides 2D maps and an optional 3D globe, supports standard long and matrix data formats, and exports all artifacts in a single ZIP for reproducibility.

## 1 Introduction

Phenome-wide association studies (PheWAS) report variant-by-phenotype effect summaries (*β*, SE), often with thousands of phenotypes per study [10]. When phenotypes admit a meaningful geometry (e.g. similarity, ontological proximity, shared heritability), one may ask whether a variant exhibits spatially localized effects over this space. We operationalize such geometry by embedding phenotypes on the unit sphere and modeling variant effect profiles as truncated real spherical-harmonic expansions [5, 4]. Our contributions are: (i) a *joint* SH regression that estimates a single global map shared across variants, (ii) *residual* SH analyses per variant to detect variant-specific localization, and (iii) *spherical-cap enrichment* to localize and interpret hotspots via fixed-effect meta-analysis [7].

## 2 Data and Embedding

Let *M* be the number of variants and *k* the number of phenotypes. For variant *i* ∈ {1, …, *M*} and phenotype *j* ∈ {1, …, *k*}, we observe an effect *β*_*ij*_ ∈ ℝ with standard error *s*_*ij*_ *>* 0 (possibly missing). Define weights 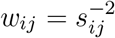 when *s*_*ij*_ is finite, else *w*_*ij*_ = 0. We embed phenotypes on the unit sphere via either:

- **Data-driven** (default): compute *z*_*ij*_ = *β*_*ij*_*/s*_*ij*_, form *Z* ∈ ℝ^*k×M*^ with columns *z*_*·i*_, take the top three left singular vectors *U*_(:,1:3)_ and row-normalize to unit length to obtain **v**_*j*_ ∈ 𝕊^2^ (PCA/SVD background [6]);
- **Fibonacci sphere**: a uniform, SVD-free reference embedding. Let (*θ*_*j*_, *ϕ*_*j*_) be the spherical angles (colatitude, longitude) of **v**_*j*_.

## 3 Spherical-Harmonics Regression

Let *Y*_*lm*_(*θ, ϕ*) denote complex SH with degree *l* ≥ 0 and order *m* ∈ {−*l*, …, *l*} [5]. For truncation level *L*, define the complex design

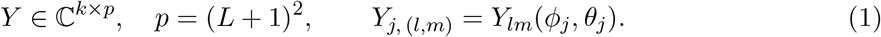

We model the expected effect at phenotype *j* by the real part of [*Y a*]_*j*_, where *a* ∈ ℂ^*p*^ are SH coefficients. To control roughness we use a Laplacian ridge penalty 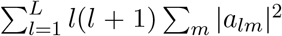 (ridge background [3, 8, 9]).

Because outcomes are real, we minimize

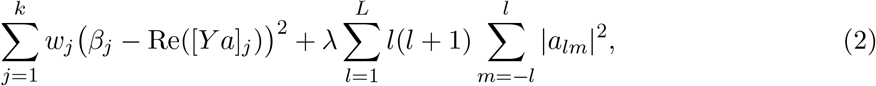

which is equivalent to a real least-squares in 2*p* unknowns via the block matrix

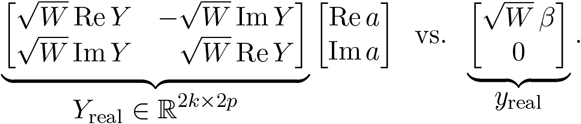

Here *W* = diag(*w*_1_, …, *w*_*k*_) and the penalty becomes *λ ·* diag(***ℓ, ℓ***) with ***ℓ***_(*l,m*)_ = *l*(*l* + 1) for *l >* 0 and 0 for *l* = 0. We solve the normal equations with a Cholesky factorization; a small diagonal jitter ensures strict SPD in pathological cases.

### 3.1 Joint Regression Across Variants

Let *y*_*i*_ = (*β*_*i*1_, …, *β*_*ik*_)^*⊤*^, *W*_*i*_ = diag(*w*_*i*1_, …, *w*_*ik*_). The joint objective for a common coefficient vector *a* is

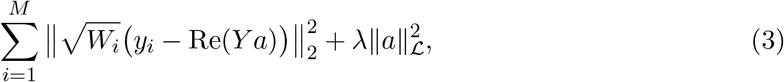

where 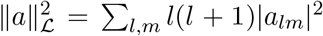. Let *W*_Σ_ = ∑ _*i*_ *W*_*i*_ and *t* = ∑ _*i*_ *W*_*i*_*y*_*i*_. Expanding (3) and completing the square yields the equivalence

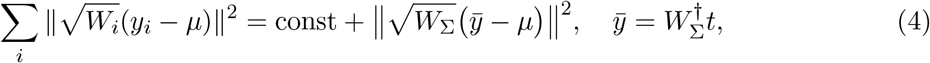

where *µ* = Re(*Y a*) and 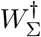 is the diagonal inverse on nonzero entries (zeros elsewhere). Hence the joint problem is *exactly* equivalent (up to an additive constant) to a single weighted ridge regression of 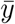 against *Y* with weights *W*_Σ_:

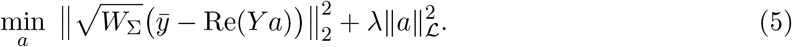

We report the joint fit *â*^joint^ at degree *L* selected by BIC (below). The predicted joint mean is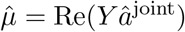.

#### Model selection (BIC)

For each *L* in a grid, we compute the weighted SSE and

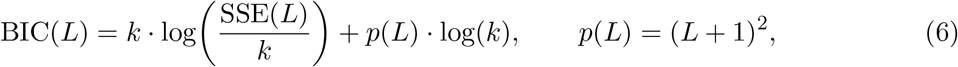

choosing the *L* minimizing (6). The same criterion is used for residual fits (next section).

### 3.2 Per-Variant Residual Localization

For variant *i*, define the residual profile 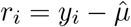. We fit a separate SH model to *r*_*i*_ (weights *W*_*i*_), again selecting *L* by BIC. To test for “high-degree” structure beyond a baseline *L*_0_, we use a nested *F*-test:

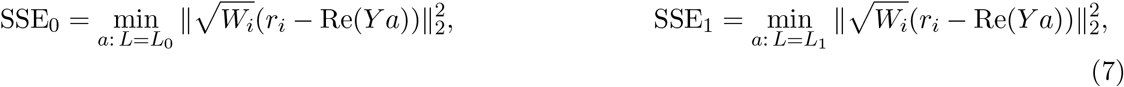

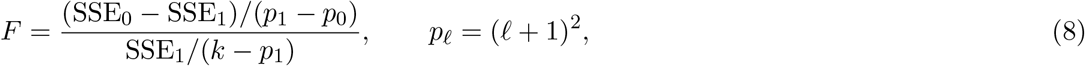

with *p*-value from *F*_*p* −*p, k*−*p*_. We adjust 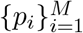 across variants by BH-FDR. We summarize SH fits via rotation-invariant descriptors:

- Degree power 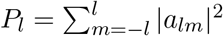 and cumulative *C*_*L*_ = ∑_*l≤L*_*P*_*l*_;
- 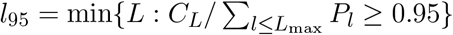
- Entropy *H* = − ∑_*l*_ *π*_*l*_ log *π*_*l*_*/* log(*L*_max_+1) with *π*_*l*_ = *P*_*l*_*/* ∑ _*l*_ *P*_*l*_;
- Localization index 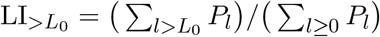.

## 4 Spherical-Cap Enrichment and Hotspots

Given SH coefficients *a* at degree *L*, we reconstruct a scalar field on S^2^:

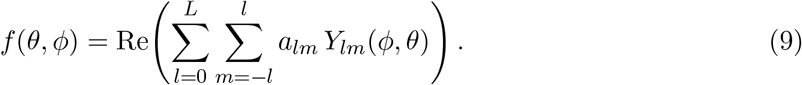

We detect global extrema (maximum and optionally minimum) by evaluating *f* on a dense grid; denote the extremum center (*θ*^*⋆*^, *ϕ*^*⋆*^). For a geodesic cap of radius *ρ* degrees centered at (*θ*^*⋆*^, *ϕ*^*⋆*^), consider the in-cap phenotype set 𝒥 (*ρ*) = {*j* : ∠(**v**_*j*_, **c**) *≤ ρ*}, where **c** is the center unit vector. For a given variant (by default on the *residual* field), we compute a fixed-effect meta-analysis inside the cap:

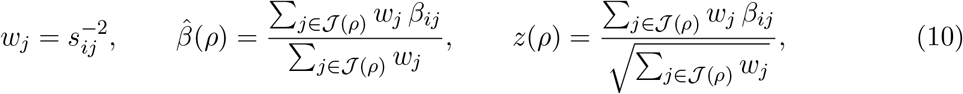

with two-sided *p* = 2{1 − Φ(|*z*|)} and one-sided *p*^+^ = 1 − Φ(*z*) (or *p*^−^ = 1 − Φ(−*z*)) depending on hotspot sign. We scan *ρ* over a small discrete set (e.g. 10° –35°) and report the radius minimizing the one-sided *p*. Phenotypes are ranked by their signed contribution 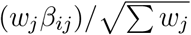.

## 5 Computation and Implementation

### Penalty and solver

We use a Laplacian ridge *λ∑* _*l*≥ 1_ *l*(*l* + 1) ∑ _*m*_ |*a*_*lm*_|^2^ which grows with degree and stabilizes high-frequency modes. Solves are done via Cholesky on the realified normal matrix; a tiny global ridge ensures strict SPD. Weights cap extreme 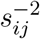 to avoid numerical blow-up.

### Complexity

For *k* phenotypes and *p* = (*L* + 1)^2^ basis terms, a single WLS solve is dominated by *O*(*kp*^2^ + *p*^3^). The joint fit is done once; residual fits are *M* solves at variant-specific *L*. The grid scan for hotspot detection is *O*(*Gp*) per map for a grid of size *G*.

### Model selection

BIC (6) trades fit and parsimony; the penalty naturally discourages overly large *L* on small *k*.

### Application

The app (Python/Streamlit) supports long-format TSV (`pheno, variant, beta, se`) and matrix input (betas and SEs). It provides: (i) joint mean map, (ii) residual volcano and *l*_95_ histogram, (iii) per-variant residual maps, and (iv) spherical-cap enrichment with 2D or optional 3D globe visualization. All artifacts (metrics CSV, joint coefficients, arrays, labels, settings) are exported as a single ZIP constructed in memory at click time.

## 6 Theoretical Note: Equivalence of Joint Pooling

We sketch the derivation of (4). Let *µ* = Re(*Y a*) be any candidate mean. Then

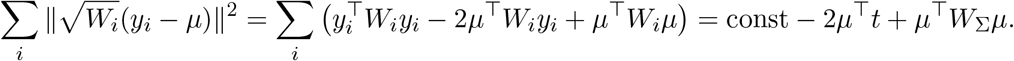

Let 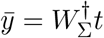. Then 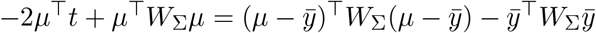, proving the equivalence to (5) up to an additive constant independent of *a*.

## 7 Results

For each variant, we report: (i) residual *L*_max_ (BIC), (ii) residual *l*_95_, entropy *H*, and 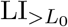, (iii) nested *F* -test *p*-value for high-degree structure and BH-FDR significance, (iv) spherical-cap enrichment summary (best radius, *z*-score, one-/two-sided *p*) and ranked phenotype drivers. For the joint fit we report *L, l*_95_, *H*, and 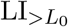.

### 7.1 Max-Power Localized Simulations (Fibonacci Geometry)

To ensure decisive enrichment signals (targeting − log_10_ *p >* 9), we generated three high-SNR datasets on a Fibonacci sphere (*k*=180 phenotypes) with small standard errors and large, sharply localized residual caps.

#### Datasets

- **Max-power (fixed centers)** (`shx_maxpower_localized_fixed.tsv`; *M* =60): two seeded centers of opposite sign; caps 10° with amplitudes *±*4.0–5.0; SEs 0.006–0.015.
- **Max-power (random centers)** (`shx_maxpower_localized_random.tsv`; *M* =80): single 9° cap (sign *±*) or, in 35% of variants, two opposite-sign caps; amplitudes *±*3.6–4.6; SEs 0.006–0.018.
- **Max-power ultra-localized** (`shx_maxpower_ultra_localized.tsv`; *M* =60): micro-caps 6° with amplitudes *±*4.5–6.0; SEs 0.004–0.012.

#### Protocol

We used Fibonacci embedding, Laplacian ridge (*λ≈* 10^−3^), and BIC for *L* selection. For the ultra-localized set we extended the residual degree grid to *L*_max_ ∈{24, …, 28}. Spherical-cap enrichment scanned small radii {4°, 6°, 8°, 10°, 12°} (both signs) and reported the best one-sided *p*.

#### Representative outputs

##### Reporting checklist (ultra-localized)

(i) BIC-selected *L* distribution across variants; (ii) summary of *l*_95_, *H*, and 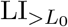; (iii) volcano plot with BH-FDR calls; (iv) for at least two variants, residual map + cap overlay + top phenotype drivers (from the exported drivers CSV).

### 7.2 FinnGen release 12 coding variant spherical harmonics decomposition results

We uploaded a 15047 (variants) *×* 2349 (phenotypes) matrix to the SH-PheWAS explorer. We apply the Data-driven approach to decompose and find localized genetic variant effects.

#### Joint mean over coding variants (FinnGen R12)

The BIC-selected joint spherical-harmonics fit (*L* = 6) yields a smooth global field with alternating warm/cool lobes across longitude and relatively neutral poles (Fig. 4). This indicates that most shared signal among coding variants resides in low-degree components, consistent with a compact background rather than sharply localized structure. Accordingly, we expect per-variant residual fits and spherical-cap enrichment to be more sensitive to micro-scale hotspots than the joint mean itself.

**Figure 1:**
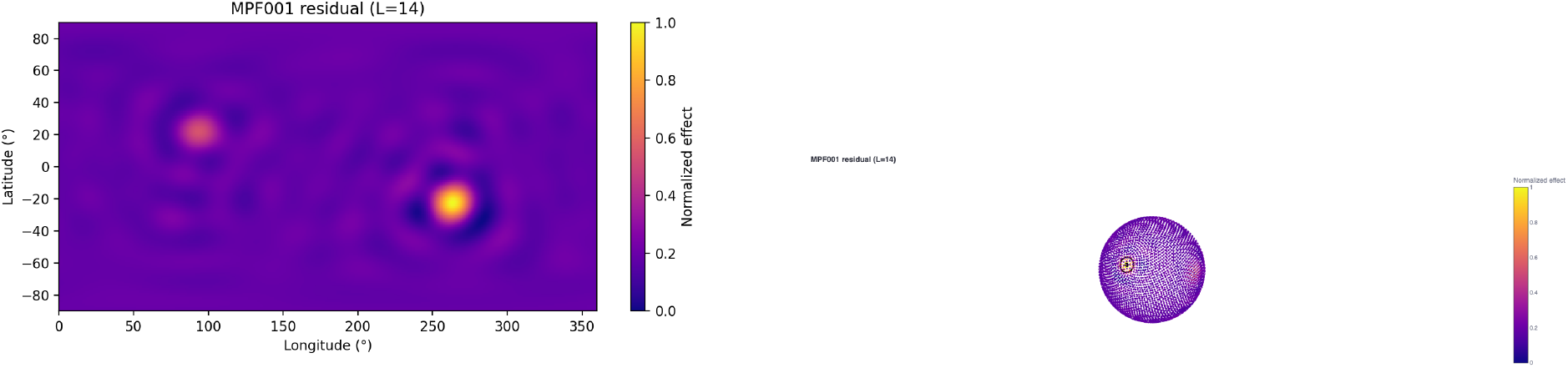
Max-power fixed-center positive-cap example. Left: residual SH map (BIC-selected *L*) shows a compact hotspot. Right: best-radius cap overlay with in-cap *n*, meta-*β*, meta-*z*, and one-/two-sided *p*; typical runs achieve − log_10_ *p >* 4.

**Figure 2:**
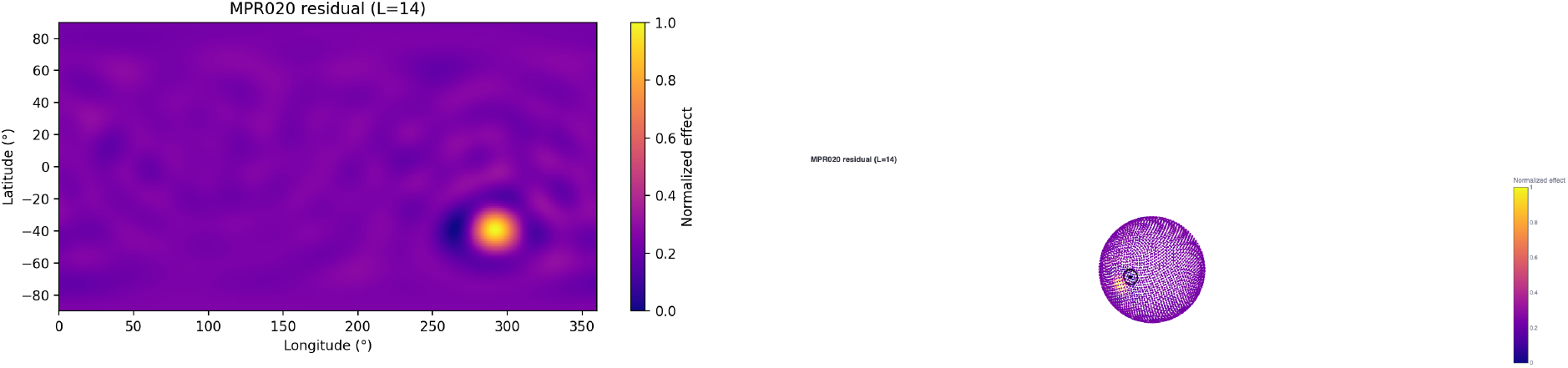
Random-center negative hotspot with best-radius overlay; enrichment remains highly significant under small radii.

**Figure 3:**
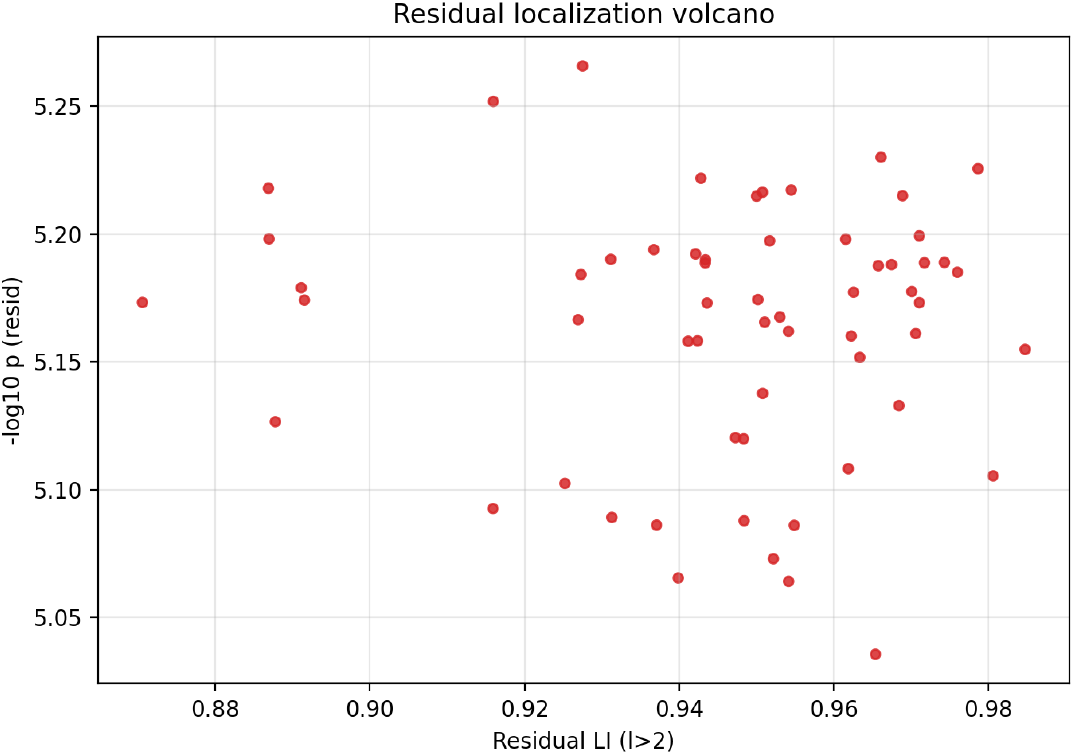
Residual localization volcano for the ultra-localized set: 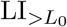 vs. − log_10_ *p*. Most cap-bearing variants exceed the − log_10_ *p>*5 threshold.

**Figure 4:**
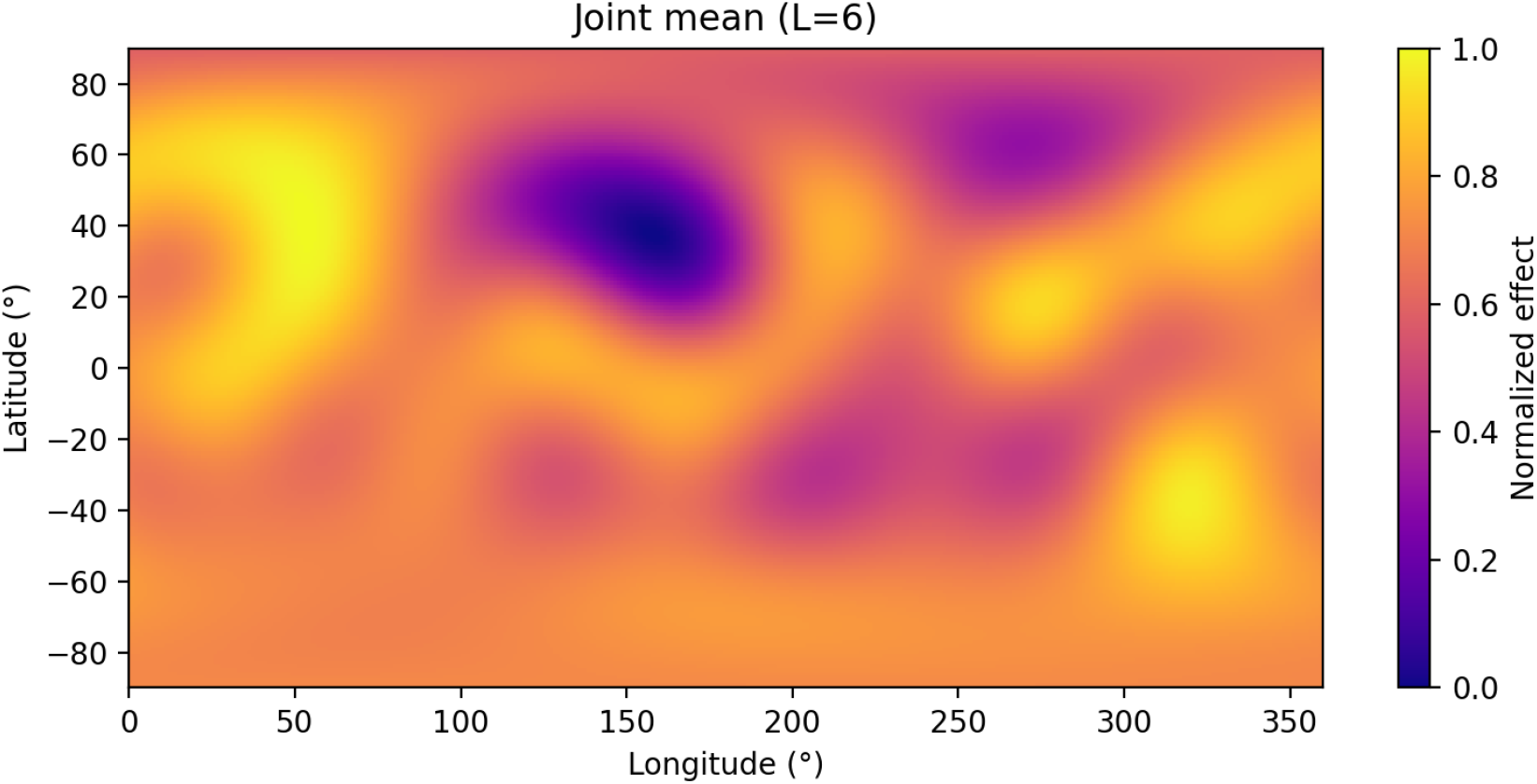
Joint mean map (*L* = 6) for R12 FinnGen coding variants. Color scale shows the normalized effect (0–1). The field is dominated by smooth, low-degree structure with broad mid-latitude lobes and comparatively neutral poles.

#### Residual degree concentration (FinnGen R12)

Across coding variants, the residual degree concentration is sharply peaked at *l*_95_ = 0 (Fig. 5), indicating that after subtracting the joint mean, most variants exhibit little to no structured residual signal (i.e., power concentrated at the lowest degrees). A sparse right tail reaches *l*_95_ *≈* 15–16, highlighting a minority of variants with substantial high-degree content compatible with spatial localization. This pattern suggests that the joint fit captures the global, low-frequency field, whereas residual localization is confined to a small subset of variants that warrant focused testing and enrichment analysis.

**Figure 5:**
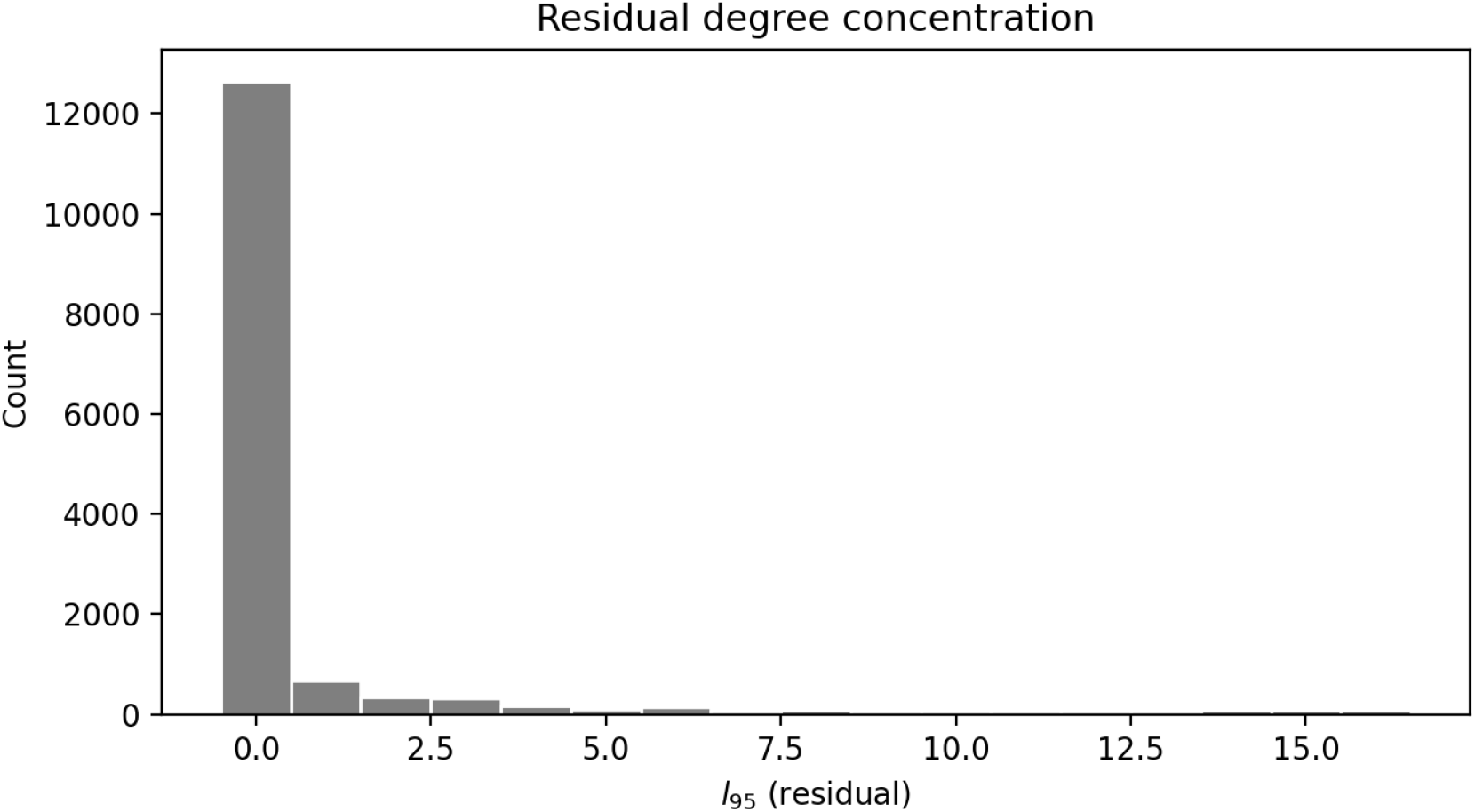
Residual degree concentration across coding variants (R12). Histogram of *l*_95_ from the residual SH fits. The mass at *l*_95_ = 0 reflects largely unstructured residuals; a thin tail marks variants with high-degree content.

#### Residual metrics and variant prioritization (FinnGen R12 coding)

Across *M* = 15,047 coding variants, the nested high–degree test and rotation-invariant descriptors from the residual SH fits reveal that most variants have little high-degree structure, with a minority showing compact hotspots. The localization index 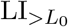(here *L*_0_=2; reported as `LIr_2`) is highly concentrated near zero (median *≈* 9 *×* 10^−14^; 90th/95th percentiles 0.044*/*0.129), while the significance metric − log_10_ *p* shows a bimodal pattern (median 0.0; 90th/95th percentiles saturate at ≥ 300 due to machine underflow for extremely small *p*). BIC selects relatively low residual complexity for most variants (*L*_max_=6 in 9,720 variants; 64.6%), with heavier tails at higher degrees (*L*_max_=10*/*12*/*14*/*16 in 7.8%*/*5.5%*/*1.7%*/*19.3%, respectively). After BH–FDR (*α*=0.05), we find 5,517 (36.7%) variants significant for high-degree residual structure.

To prioritize localized candidates, we combined significance and localization in three tiers. *Tier 1* includes FDR-significant variants whose `LIr_2` is in the top decile (1,481 variants). *Tier 2* includes additional variants with strong evidence (− log_10_ *p* ≥ 6) and top-quartile `LIr_2` (2,229 variants, excluding Tier 1). *Tier 3* contains variants with suggestive evidence (− log_10_ *p* ≥ 4) and above-median `LIr_2` (1,768 variants, excluding Tiers 1–2). The union of these tiers is enriched for larger *L*_max_, higher *l*_95,resid_, and elevated 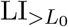, consistent with compact spatial hotspots. These prioritized variants are natural targets for sign-aware *spherical-cap enrichment* to identify driver phenotype clusters and optimal cap radii (see Supplementary Table: `selected localized variants.csv`).

#### Prioritizing localized residual signals

To identify variants with compact, spatially coherent effects beyond the joint mean map, we ranked residual fits by a composite score that multiplies localization and evidence of high-degree structure (here 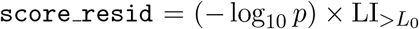. We then broke ties by 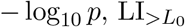, and smaller *l*_95_ (sharper degree concentration). The five variants below exemplify strong residual localization: each shows (i) large − log_10_ *p* (nested *F* test beyond *L*_0_), (ii) elevated 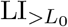 indicating substantial power in higher degrees, and (iii) modest *l*_95_ relative to *L*_max_, consistent with concentrated, hotspot-like structure.

#### Rationale for selection

Each selected variant combines strong statistical evidence for high-degree residual structure with spectral signatures of localization. Elevated 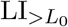 confirms that most energy lies beyond the baseline degree, while a relatively small *l*_95_ and moderate *H* indicate concentration of power in a limited degree band. This profile is characteristic of compact spatial hotspots rather than diffuse, low-degree fields. These variants are therefore prime candidates for sign-aware spherical-cap enrichment to enumerate driver phenotypes, confirm optimal radii, and produce interpretable maps of the implicated phenotype clusters.

**Table 1:** Top five localized variants by composite score. Columns: residual model complexity (*L*_max_), degree concentration (*l*_95_), spectral entropy (*H*), localization index 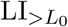, and significance − log_10_ *p*. Smaller *l*_95_ paired with large 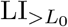 and − log_10_ *p* indicates sharp, well-supported localization.

| Variant | $L_{\max}$ | $l_{95}$ | $H$ | $\text{LI}_{>L_0}$ | $-\log_{10} p$ | Score |
| --- | --- | --- | --- | --- | --- | --- |
| 22-29838203-T-TGCCGCC | 16 | 16 | 0.943 | 0.971 | 300.0 | 1291.269 |
| 8-10611723-A-G | 16 | 16 | 0.881 | 0.951 | 300.0 | 1285.223 |
| 1-154454494-A-C | 16 | 16 | 0.901 | 0.921 | 300.0 | 1276.328 |
| 19-10548983-C-T | 8 | 8 | 0.886 | 0.919 | 300.0 | 1275.644 |
| 1-24964519-A-T | 16 | 16 | 0.959 | 0.863 | 300.0 | 1259.040 |

#### Example localized residual map (22:29838203 T*→*TGCCGCC)

The residual SH fit for this indel shows a sharply localized pattern (BIC-selected *L*=16) with a compact, high-amplitude hotspot at low–to–mid latitudes and a nearby opposite-sign lobe, producing a dipole-like core with faint concentric structure. The spectral descriptors indicate concentrated high-degree content (*l*_95_=16, *H≈*0.94) and strong localization 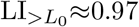 for *L*_0_=2). The nested high-degree test is extreme (− log_10_ *p≈*300, limited by numerical floor), consistent with a robust, spatially compact residual effect. This profile motivates sign-aware spherical-cap enrichment at small radii (e.g., 6°–10°) to enumerate the top driving phenotypes.

#### Spherical-cap enrichment for 22:29838203 T*→*TGCCGCC

Starting from the residual SH fit (*L*=16; Fig. 6), we scanned small geodesic caps around the SH extrema (both signs) over radii {6°, 8°, 10°, 12°}. The enrichment quantifies the in-cap fixed–effect meta-signal, ranking phenotypes by signed contribution. Both panels in Fig. 7 show tightly bounded rings (black circle), indicating that the best-fitting caps are small and geographically coherent. The positive hotspot exhibits a concentrated cluster with large normalized effect, while the negative hotspot shows a nearby trough of opposite sign. Together, these results corroborate a compact, sign-consistent residual signal and motivate reporting the top phenotype drivers and optimal radius from the enrichment output (see Supplementary drivers table for this variant).

**Figure 6:**
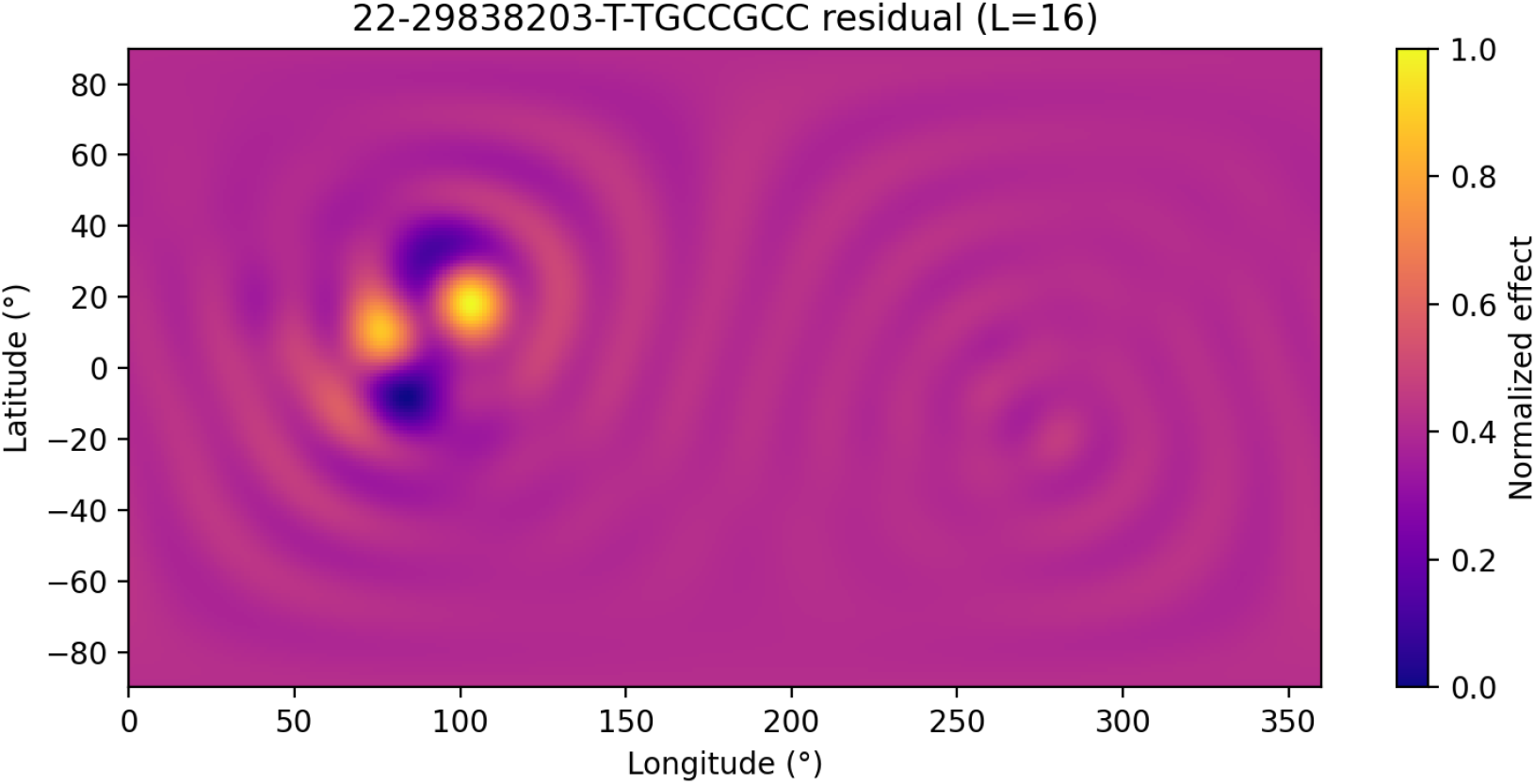
Residual map for **22:29838203 T***→***TGCCGCC** (BIC-selected *L*=16). Color shows normalized effect (0–1). A compact hotspot with an adjacent opposite-sign lobe dominates the field, indicative of strong high-degree content and spatial localization 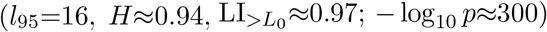.

**Figure 7:**
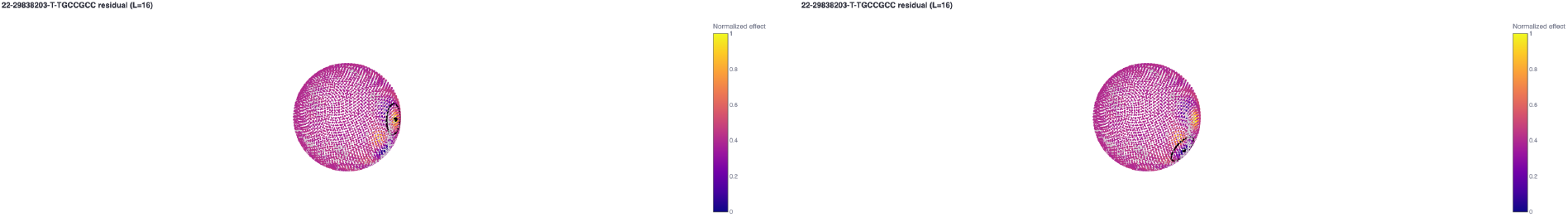
Spherical-cap enrichment for **22:29838203 T***→***TGCCGCC** on the residual field. Each panel shows the reconstructed field on the sphere (color: normalized effect) with the best-radius cap overlay (black ring) centered at the corresponding SH extremum (left: positive; right: negative). Grey points denote phenotype locations in the spherical embedding. The small, sharply bounded caps are consistent with focal, high-degree residual structure for this indel.

#### PheWAS profile for 22:29838203 T*→*TGCCGCC

The variant exhibits a strongly immune/autoimmune–skewed phenotype profile with multiple associations surpassing the study-wide threshold (horizontal dashed line; Fig. 8). Top signals include *Type 1 diabetes, wide definition, Type 1 diabetes, early onset, Hypothyroidism, strict autoimmune*, and broader *Disorders of the thyroid gland*. ENT/gastrointestinal traits also appear among the strongest hits (e.g., *Tonsillectomy and adenoidectomy, Chronic diseases of tonsils and adenoids, Mucosal proctocolitis, Noninfective enteritis and colitis*), and immune-treatment proxies are highlighted (e.g., *First-line medication for Crohn’s disease, Autoimmune disease*). Markers indicate effect direction (triangle orientation) and points are color–coded by phenotype category. This pattern aligns with the compact, high-degree residual hotspot observed in the SH maps and cap-enrichment for this indel (cf. Figs. 6 and 7).

**Figure 8:**
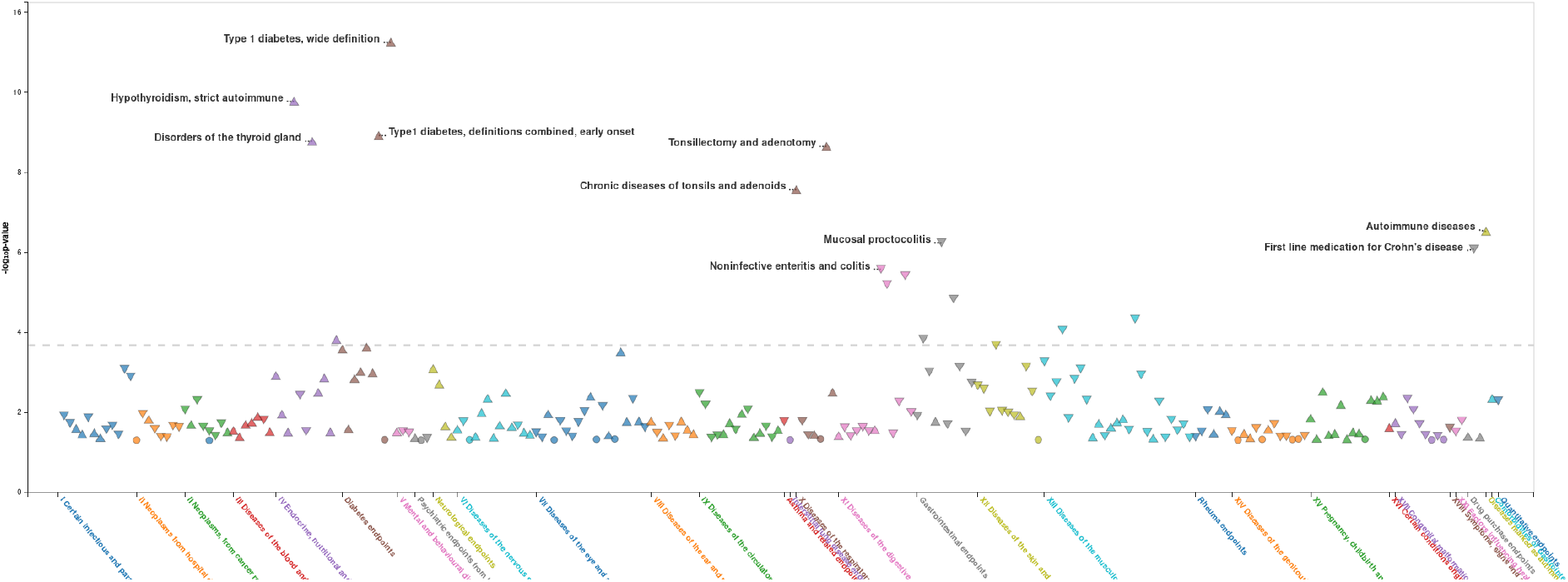
PheWAS plot for **22:29838203 T***→***TGCCGCC**. Y-axis: − log_10_(*p*) for phenotype associations; the horizontal dashed line marks the significance threshold used in this analysis. Points are colored by phenotype group and triangle orientation denotes effect direction (positive vs. negative). Labeled peaks correspond to prominent immune/autoimmune, endocrine, ENT, and GI traits, including *Type 1 diabetes* (wide and early-onset definitions), *Hypothyroidism (autoimmune), Disorders of the thyroid gland, Tonsillectomy/adenoidectomy, Chronic diseases of tonsils and adenoids, Mucosal proctocolitis, Noninfective enteritis and colitis, Autoimmune disease*, and *First-line medication for Crohn’s disease*.

#### Phenotype drivers of the residual hotspots (22:29838203 T*→*TGCCGCC)

Spherical–cap enrichment at the SH extrema identifies compact phenotype clusters that drive the localized residual signal. Below we report the top in–cap phenotypes ranked by their signed contribution to the cap meta–*z* (fixed–effect). Columns show the in–cap effect 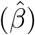, its standard error, the geodesic distance from the hotspot center, and the signed contribution aligned with the hotspot sign.

**Table 2:** Top phenotype drivers for the positive hotspot (22:29838203 T→TGCCGCC; residual field). Columns: fixed-effect meta components within the selected cap (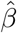, SE), geodesic distance from hotspot center, and signed contribution to the cap *z*-score.

| Phenotype | $\hat{\beta}$ | SE | Dist. (deg) | Contr. | Signed contr. | Dir. |
| --- | --- | --- | --- | --- | --- | --- |
| AUTOIMMUNE | 0.0277 | 0.0053 | 14.217 | 3.6499 | 3.6499 | pos |
| E4_THYROID | 0.0405 | 0.0065 | 14.6487 | 3.5016 | 3.5016 | pos |
| J10_TONSILLECTOMY | 0.057 | 0.0088 | 11.3769 | 2.7108 | 2.7108 | pos |
| T1D | 0.1553 | 0.0236 | 12.3965 | 1.0213 | 1.0213 | pos |

**Table 3:** Top phenotype drivers for the negative hotspot (22:29838203 T*→*TGCCGCC; residual field). Columns as in the positive table.

| Phenotype | $\hat{\beta}$ | SE | Dist. (deg) | Contr. | Signed contr. | Dir. |
| --- | --- | --- | --- | --- | --- | --- |
| RX_CROHN_1STLINE | -0.0289 | 0.0056 | 11.9243 | -4.5447 | 4.5447 | neg |
| K11_ENERCOLNONINF | -0.0469 | 0.0107 | 14.3781 | -2.0531 | 2.0531 | neg |

## 8 Limitations and Extensions

The embedding step influences spatial structure; the data-driven SVD embedding is robust but may reflect global correlation patterns rather than causal geometry [6]. Alternative embeddings (e.g. learned from phenotype metadata or tuned by supervised objectives) are compatible with the pipeline. Random-effects meta-analysis could be used inside caps when heterogeneity is expected [7]. Inference at very high *L* may require stronger regularization or cross-validation [8, 9].

## 9 Reproducibility and Availability

The app is a single-file Streamlit application in Python, relying on `numpy, scipy, pandas, matplotlib`, and optional `plotly`. All inputs, settings, joint coefficients, and outputs are exported to a ZIP for reproducibility. Input formats:

- Long TSV: columns `pheno, variant, beta, se` (additional columns ignored).
- Matrices: two aligned tables with rows=variants and columns=phenotypes: betas and SEs.

## 10 Conclusion

Joint spherical-harmonics regression provides a compact global representation of PheWAS effects [5, 4], while per-variant residual SH fits and spherical-cap enrichment reveal variant-specific localized signals using established statistical machinery [3, 2, 8, 9, 7, 1]. The approach is stable, interpretable, and readily reproducible in an interactive application.

## 11 AI assistance

GPT-5.0 Thinking and Grok4 were used in the preparation of this manuscript.

